# Paternal age affects offspring’s behavior possibly via an epigenetic mechanism recruiting a transcriptional repressor REST

**DOI:** 10.1101/550095

**Authors:** Kaichi Yoshizaki, Tasuku Koike, Ryuichi Kimura, Takako Kikkawa, Shinya Oki, Kohei Koike, Kentaro Mochizuki, Hitoshi Inada, Hisato Kobayashi, Yasuhisa Matsui, Tomohiro Kono, Noriko Osumi

## Abstract

Advanced paternal age has deleterious effects on mental health of next generation. Using a mouse model, we have confirmed that offspring derived from aged fathers showed impairments in behavior and abnormalities in the brain structure and activity. Comprehensive target DNA methylome analyses revealed in aged sperm more hypo-methylated genomic regions, in which REST/NRSF binding motif was enriched. Gene set enrichment analyses also identified enrichment of “REST/NRSF target genes”, in addition to “Late-fetal genes” and autism spectrum disorder-related “SFARI genes”, in up-regulated genes of developing brains from aged father. Indeed, gene sets near hypo-methylated genomic regions with REST/NRSF binding motif were also enriched in up-regulated genes of developing brains. Taken altogether, DNA hypo-methylation due to paternal aging in sperm will induce leaky expression of REST/NRSF target genes in the developing brain, thereby causing neuronal abnormalities and subsequent behavioral alteration in offspring.

## Introduction

Human epidemiological studies have indicated that advanced paternal age is related to higher risks for various psychiatric disorders, such as schizophrenia ^1^, autism spectrum disorders (ASD) ^2, 3, 4^, early onset of bipolar disorder ^5^, reduced IQ ^6^, and impaired social functioning in adolescents ^7^ in their offspring. In rodents, paternal aging actually induces offspring’s abnormal behaviors such as learning deficit, impaired social behavior, and hyper anxiety ^8, 9, 10, 11^. However, molecular mechanisms underlying the transgenerational inheritance have not been fully understood to date.

A previous exome analysis for ASD revealed that there are more paternally-derived *do novo* mutations ^12, 13^, although there is no direct evidence relating these mutations with mental diseases. Epigenetic changes are also involved in sperm aging. DNA methylation levels in human sperm show age-related changes ^14^. A longitudinal analysis of ASD-high risk children has indeed showed correlation between DNA methylation in father’s sperm and risk for ASD in their children ^15^. This is partially modeled in mice; Milekic and his colleagues have performed methylation Mapping Analysis by Paired-end Sequencing (methyl-MAPS) and revealed significant hypo-methylation in old sperm as well as offspring’s brain ^11^. However, it is still largely unknown how global changes in the DNA methylation alter potential dysregulation of gene expression within the offspring’s brain and their behavior.

In the present study, we first evaluated in mice influence of paternal aging by selected behavior tasks related with neurodevelopmental disorders. We confirmed that advanced paternal age affected body weight, maternal separation-induced vocal communication (USV), sensorimotor gating and spatial learning in F1 offspring. Structural and neuronal abnormalities were observed in the brain regions related with behavior impairment. Target methylome analyses using young and aged sperm identified genome-wide differentially methylated regions (DMRs) including a unique consensus motif of binding sites for REST/NRSF, a transcriptional repressor important for neural differentiation ^16, 17^. Expression profiling of the developing brain suggested potential involvement of REST/NRSF target genes and precocious neurogenesis due to paternal aging. These results provide a scenario that abnormal sperm methylation in the aged father results in leaky dysregulation of REST/NRSF-target genes critical for proper development of the brain, thereby affecting offspring’s behavior.

## Results

### Advanced paternal age affected body weight, vocal communication, sensorimotor gating and spatial learning in F1 offspring

To examine transgenerational influence of paternal aging on offspring behavior, we performed selected behavior tests used for mouse models of neurodevelopmental disorders (Fig.S1) ^18^. Wild type (C57BL/6J) F1 offspring mice were obtained from young (3 month) or aged (>12 month) male mice mated with young (3 month) virgin female mice (Fig.1a).

**Figure 1.**
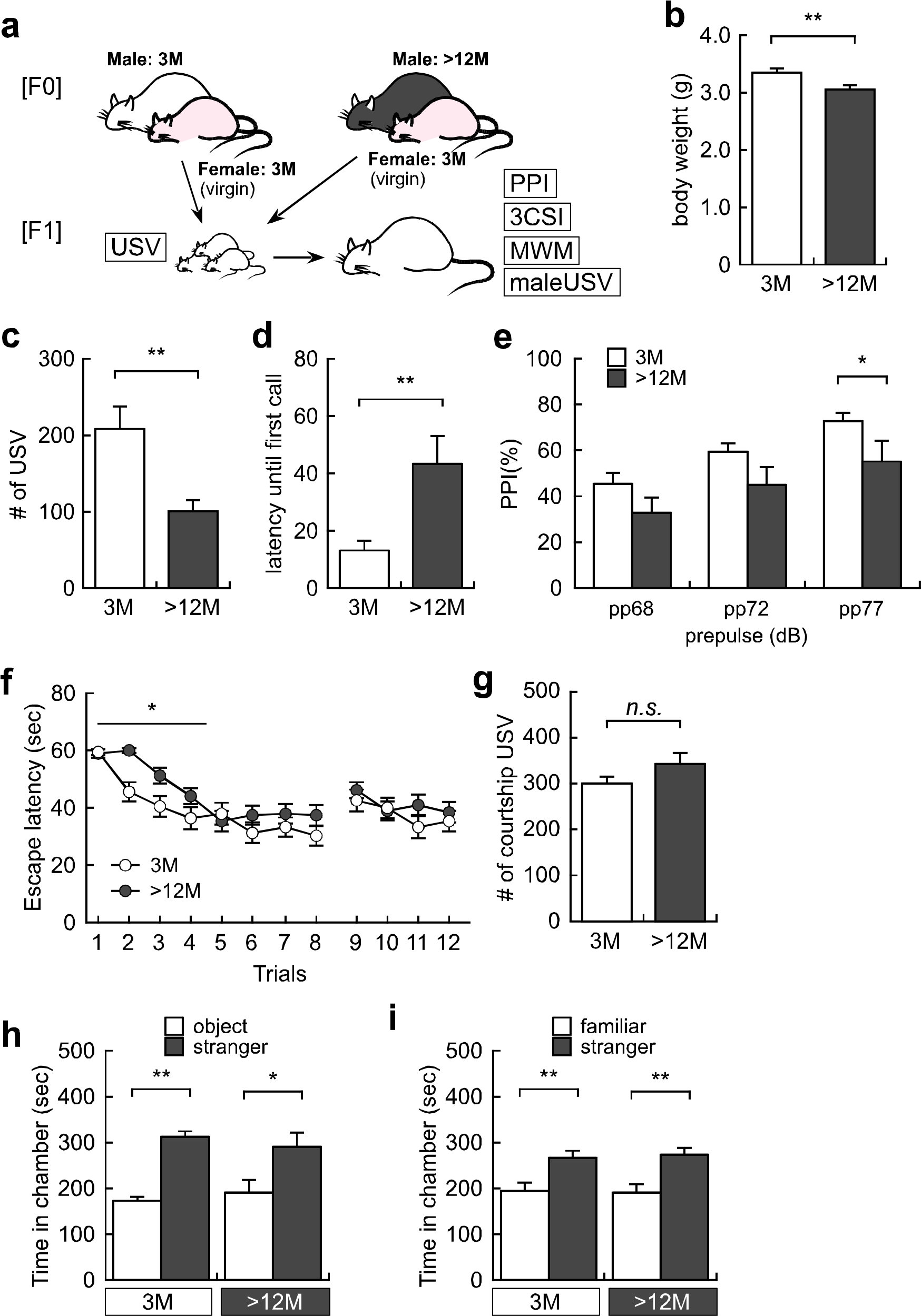
Advanced paternal age affected body weight and behavior phenotypes in F1 offspring derived from aged father. (a) Experimental design of F1 offspring derived from either young or aged father. (b) Body weight, (c) the number of maternal separation-induced ultrasonic vocalization (USV) calls, and (d) latency until first USV calls at postnatal day 6. (e) pre-pulse inhibition score at pp68 dB, pp72 dB and pp77 dB, (f) spatial and reversal learning in Morris water maze test, (g) courtship USV to female mouse, and (h) sociability and (g) social novelty in three chamber social interaction test at month-old in F1 offspring derived from young or aged father. All data are presented by the mean ± SEM. \**p*<0.05, \*\**p*<0.01, determined by Student *t* test or two-way ANOVA followed by post hoc test with Bonferroni method.

At postnatal day 6 (P6), we measured body weight and recorded maternal separation-induced pup’s ultrasonic vocalization (USV) that reflects vocal communication in rodents ^19, 20, 21, 22^. F1 offspring pups derived from aged father exhibited significant loss of body weight by 8.6% compared with that derived from young father (3.36 ± 0.06 g: young father vs 3.07 ± 0.07 g: aged father, *p*=0.003, Fig.1b), which is consistent with previous studies in the mouse and human ^23, 24, 25^. Moreover, the number of USV in F1 pups derived from aged father was significantly decreased by 52.1% compared with that derived from young father (209.6 ± 28.0 calls: young father vs 100.5 ± 16.1 calls: aged father, *p*=0.021, Fig.1c). Latency until the first USV in F1 pups derived from aged father was correspondingly delayed than that derived from young father (18.1 ± 5.3 sec: young father vs 35.5 ± 5.5 sec: aged father, *p*=0.027, Fig.1d). These phenotypes are consistent with many other models of ASD ^26, 27, 28^.

Using adult offspring, we next performed the prepulse inhibition (PPI) test, an endophenotype for ASD and schizoophrenia ^29, 30^. The PPI score with 77 dB prepulse stimuli against 120 dB startle stimuli, but not 68 and 72 dB prepulse stimuli, was decreased by 24.1% in F1 offspring derived from aged father compared to that derived from young father (73.1 ± 3.4 %: young father vs 55.5 ± 8.9 %: aged father, Fig.1e, *p*=0.029).

In addition, we conducted the Morris water maze test to examine spatial learning and memory. F1 offspring derived from aged father showed higher scores for escape latency in the early phase of spatial learning, but not the late phase of spatial learning, meaning delay in special learning (Fig.1f). Since there was no difference in reversal learning, we consider that the offspring derived from aged father did not show a repetitive phenotype in this paradigm.

Regarding other behavior tests, both F1 offspring derived from young and aged father exhibited the comparable number of courtship USV (Fig.1g). Furthermore, there was no difference in social behavior in the 3-chamber social interaction test; both F1 offspring derived from young and aged father spent significantly much time with stranger mouse than object compartment (Fig.1h). Similarly, both of the F1 offspring showed interest with a stranger mouse than a familiar mouse (Fig.1i).

In summary, paternal aging in mice has indeed resulted in abnormalities of the behavior phenotypes (i.e., impairment in vocal communication in pups, and defects of sensorimotor gating and spatial learning in adult offspring) that can be considered to reflect some aspects of the endophenotypes of neurodevelopmental disorders such as ASD and schizophrenia.

### Advanced grand-paternal age affected body weight, but not behavior phenotypes in F2 offspring

To examine whether transgenerational influence of paternal aging was inherited to the subsequent generation, we conducted behavior analyses using F2 offspring. Young (3-month) F1 offspring mice derived from either young or aged grandfather were mated with young female mice to obtain F2 mice (Fig.S2a). We found that body weight at P6 in F2 offspring derived from aged grandfather remained decreased by 4.9% (less than the difference in F1, yet statistically different) compared to that derived from young grandfather (3.14 ± 0.05 g: young father vs 2.98 ± 0.05: aged grandfather, *p*=0.039, Fig.S2b).

In contrast, the number of USV and latency until the first USV at P6 as well as sensorimotor gating and spatial learning in adult were unchanged between F2 offspring derived from young father born to young and aged grandfather (Fig.S2c-f). Likewise, courtship USV and social behavior in adult were similar between F2 offspring derived from young or aged grandfather (Fig.S2g-i). These results suggest that transgenerational influence of paternal aging on the next generation is differential among traits; reduction of body weight can still remain in F2 generation, while impairment of vocal communication, sensorimotor gating and spatial learning can be canceled.

### Abnormal brain structures in offspring derived from aged father at postnatal day 6

We next addressed whether paternal aging impaired the offspring brain at the levels of structures and neuronal activity at postnatal day 6 (P6) when impairment of maternal separation-induced USV was observed. We noticed that cortical thickness was reduced in offspring derived from aged father (Fig.2a, b). To determine whether impaired cortical thickness was associated with specific neuronal types, we focused on the six cortical layers using layer specific markers (Fig.2c, d) ^31, 32, 33, 34^. Reduction of thickness was specifically observed in Tbr1^+^/Ctip2^−^ layer 6 neurons, but not in Satb2^+^/Ctip2^−^ layer 2-4 and Ctip2^+^ layer 5 neurons in the offspring’s brain derived from aged father (Fig.2e). Correspondingly, 39.8% reduction of the number of neurons were confirmed in Tbr1^+^/Ctip2^−^ layer 6 neurons in the offspring brain derived from aged father (Fig.2f). We also noticed that cell density in layer 6 (Tbr1^+^/Ctip2^−^) was reduced to 59.6% in the offspring brain derived from the aged father, but not in layer 5 (Ctip2^+^) nor layer 2-4 (Satb2^+^/Ctip2^−^) (Fig.2g). These findings imply that early neurogenesis producing deep layer neurons is impaired in the neocortex of the offspring derived from the aged father. Since lesions of the motor cortex in human severely impair or eliminate the ability to produce vocalization ^35, 36^, thinner motor cortex can be considered to relate with less vocal communication in pups derived from aged father.

**Figure 2.**
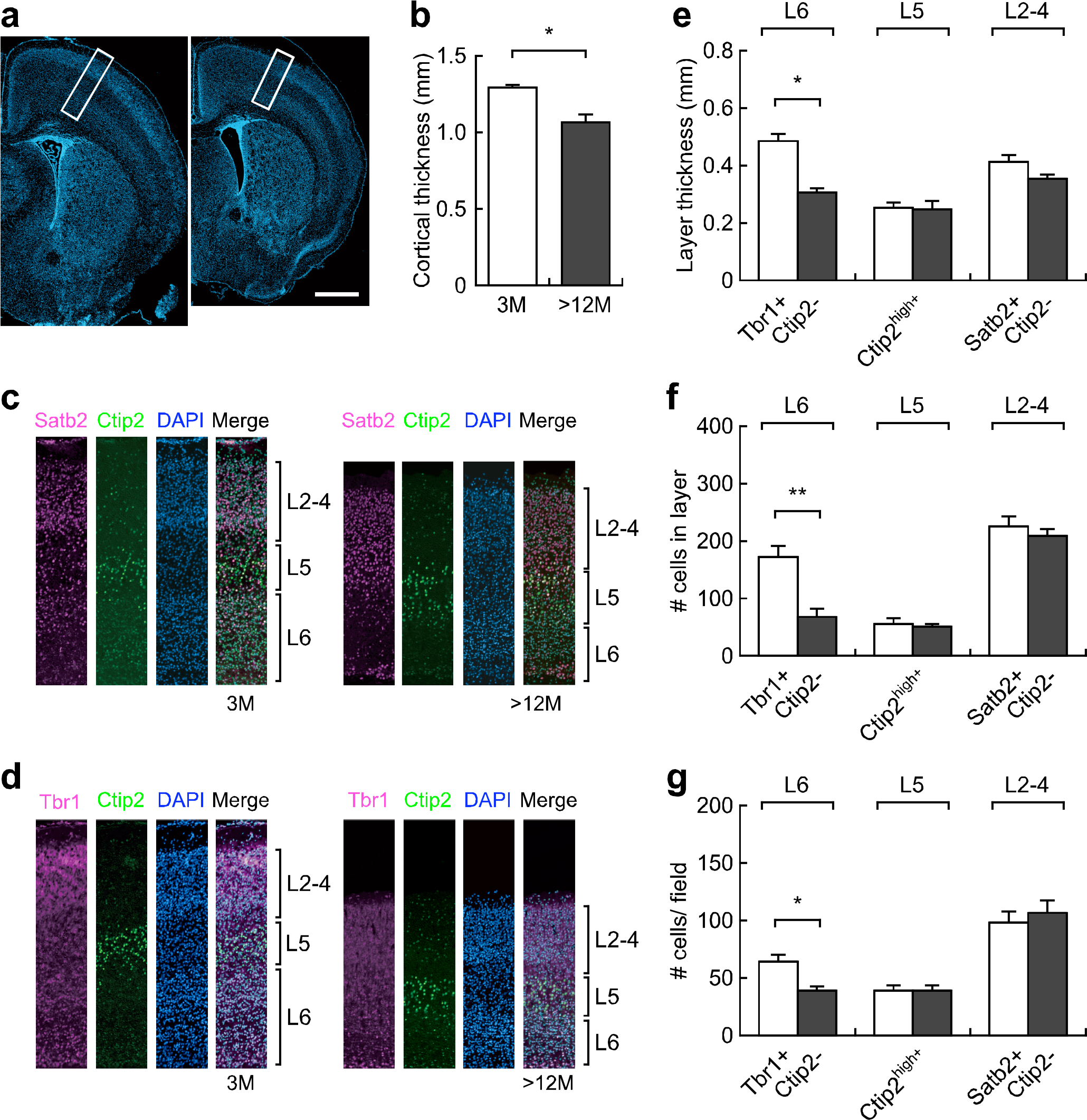
Abnormal cortical structures in the offspring brain derived from aged father at early postnatal period. (a, b) Representative image and thickness of cerebral cortex in offspring brain derived from young (left) or aged (right) father at postnatal day 6. Scale bar: 1 mm. (n=4-6 in each group). (c, d) Representative image of each cortical layer marker in offspring brain derived from young or aged father. Satb2^+^:Ctip2^−^, layer 2-4:Ctip2^high^, layer 5; Tbr1^+^:Ctip2^−^, layer 6. (e) Thickness, (f) cell number and (g) cell density in each cortical layer in offspring brain derived from young or aged father. All data are presented by the mean ± SEM. \**p*<0.05, \*\**p*<0.01, determined by Student *t* test.

### Impaired neuronal activities of anxiety-related brain regions in offspring derived from aged father at postnatal day 6

We further examined neuronal activities of the offspring at P6 when we found the behavioral impairment at the earliest stage. We focused on the paraventricular nucleus of thalamus (PVT), the basolateral amygdala (BLA), and the piriform cortex (Fig. 3a-c) because these brain regions are reported to reflect with maternal separation-induced anxiety ^37, 38^. c-Fos expression was dramatically induced 2 hour after USV measurement, particularly, in the PVT and in BLA, but not in the piriform cortex (Fig. 3d-f). The number of USV emitted by individual mouse pups showed a significant correlation with the number of c-Fos^+^ cells 2-hour after USV measurement in the PVT but not in other two brain regions (Fig. 3g-i). Interestingly, the number of c-Fos^+^ cells was dramatically reduced in the PVT and BLA, but not in the piriform cortex, after the USV measurement in offspring derived from aged father (Fig. 3j-l), which was not due to the decrease of neurons in these brain regions (Fig. 3m-o). These data clearly demonstrate that PVT is the most responsible brain regions for maternal separation-induced anxiety and paternal aging impaired offspring’s neuronal activities in anxiety-related brain regions, PVT and BLA by maternal separation at the early postnatal stage.

**Figure 3.**
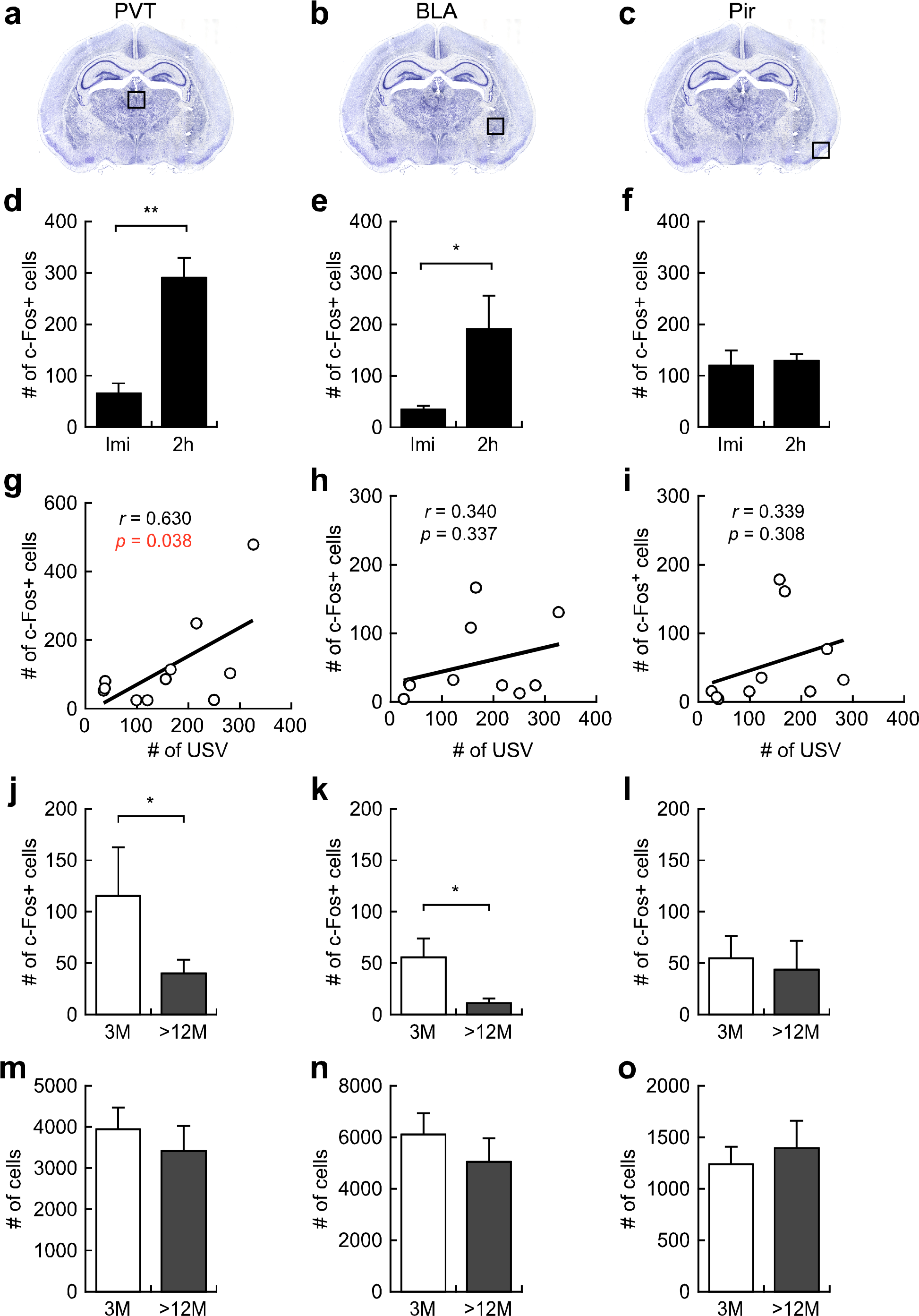
Impaired neuronal activity after maternal separation in offspring brain derived from aged father. (a-c) Brain images of anxiety-related brain regions, i.e., the paraventricular nucleus of the thalamus (PVT, a, d, g, j, m), the basolateral amygdala (BLA, b, e, h, k, n) and the piriform cortex (Pir, c, f, i, l, o) at the early postnatal stage ^37, 38^. (d-f) The number of c-Fos positive cells immediately or 2 hours after USV measurement in PVT (d), BLA (e) and Pir (f). (g-i) Correlation analyses between the number of USV calls and of c-Fos positive cells in PVT (g), BLA (h) and Pir (i). (j-l) The number of c-Fos positive cells in PVT (j), BLA (k) and Pir (l) of pups derived from young and aged father. (m-o) Total cell number in PVT (m), BLA (n) and Pir (o). All data are presented by the mean ± SEM. \**p*<0.05, \*\**p*<0.01, determined by Student *t* test.

### Comprehensive target methylome analysis identified more hypo-methylated regions than hyper-methylated regions in aged sperm

To uncover the molecular basis for transgenerational influence of paternal aging, we investigated epigenetic changes in sperm because the behavior phenotypes were canceled in F2 generation (Fig. S2). Target methylome analyses by SureSelect Methyl-Seq, Agilent Technology ^39^ (Fig. 4a) were conducted using 4 young and 9 aged sperm samples since epigenetic changes accumulate during aging (see Review by Fraga and Esteller, 2007 ^40^). Sequence analysis of the sperm samples yielded 17.5–50.0 million uniquely mapped reads (40.1%–56.3% of total reads) that could map to the mm10 reference (Fig. S3a). The average level of CpG methylation at the targeted regions was 33.3%–34.6% and 27.7%-36.2% in young and aged father data sets, respectively. The average depth of each sample was 11.3–27.6, which covered 44.2%–62.9% of the target regions at a depth of 10 (Fig. S3a). Correlation coefficients showed similarly high values—i.e., 0.98–0.99 (Fig. S3b), indicating that the obtained sequence data was sufficient for further analysis of age-specific methylation changes.

**Figure 4.**
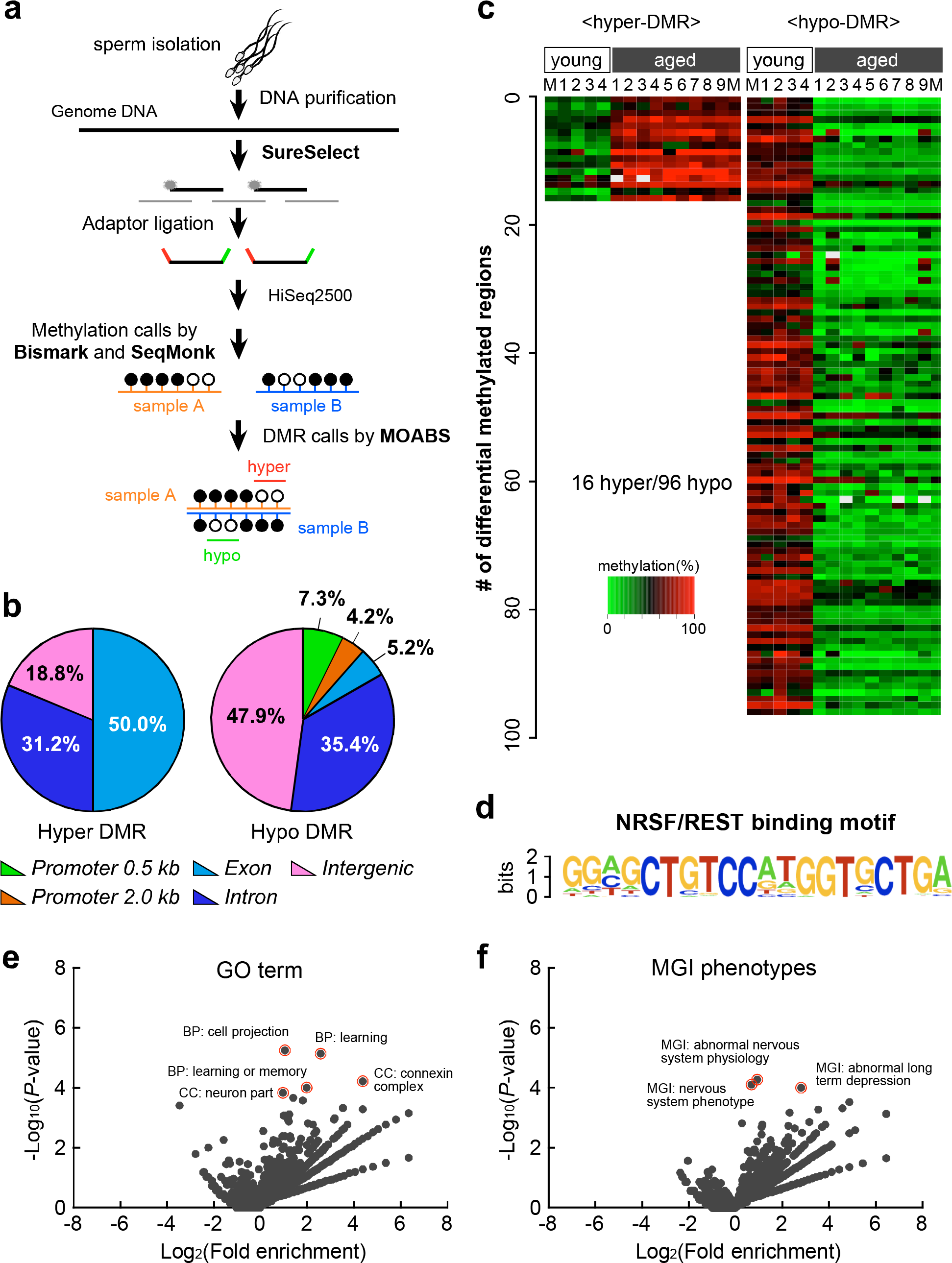
Comprehensive target methylome analysis identified more hypo-methylated genomic regions including REST/NRSF binding motifs in aged sperm. (a) Sperm from young (3-month-old, n=4) and aged (18-month-old, n=9) father was applied for target methylome analyses using the SureSelect system, PBAT library construction, and Illumina high-throughput sequencing. The successive computational steps are as follows; mapping the reads to genome (Bismark and SeqMonk), calling DMRs between two methylomes (MOABS), and scanning the DMR sequences for known motifs (HOMER). (b) Pie chart showing the distribution of DNA hyper- (left) and hypo- (right) methylated regions (promoter [0.5 kb or 2.0 kb upstream of TSS], exons, introns, intergenic regions) in length-base analyses. (d) Motif analysis of DNA hypo-methylated regions in aged sperm (19/96). (e, f) Volcano plots showing GO terms (s) and MGI phenotypes (f) enriched for genes nearby hypo-methylated regions relative to other RefSeq genes.

Although the SureSelect Methyl-Seq is designed to enrich transcription starting site (TSS) regions and putative cis-regulatory regions ^41^, our results by the length-base analysis showed the majority of our hyper-methylated genomic regions in gene-body regions (50.0% in exons and 31.2% in introns, respectively), while almost half (47.9%) of the hypo-methylated regions were detected in intergenic regions (Fig. 4b). Only 5.2% of DMRs were detected in exon regions, and 11.5% were detected in promoter regions by the length-base analysis (Fig. 4b). The count-base analysis also revealed that the majority of our hyper-methylated genomic regions in gene-body regions (31.9% in exons and 48.8% in introns, respectively), while almost half (51.3%) of the hypo-methylated regions were detected in intergenic regions, and only 4.3% of DMRs were detected in exon regions, and 3.1% were detected in promoter regions (Fig. S4). These bioinformatic findings imply that both hyper- and hypo-methylated genomic regions due to paternal aging were highly localized in exon and intron, considering the proportion in the whole genomic sequence.

Finally, we analyzed DMRs in aged sperm using the Model-based Analysis of Bisalfite Sequence data (MOABS) tool with Method2 and minDepthForComp=10^42^. Using ×10 ≧ depth CpG site data (Table. S1 and S2). These DMRs were found across the entire chromosomes except sex chromosomes. This may be because the target regions of SureSelect that capture X and Y chromosomes are extremely small in number (7,347 and 114, respectively) compared with the genome size of X and Y chromosomes (171,031,299 and 91,744,698, respectively). Interestingly, heat map analysis clearly classified the sperm data sets into young (n=4) and aged (n=9) groups with 16 hyper- and 96 hypo-methylated unique DMRs in aged mouse sperm (Fig. 4c). It is thus suggested that sperm aging induces hypo-methylation, rather than hyper-methylation, of the chromosomal DNA.

### Comprehensive target methylome analysis identified REST/NRSF binding motif in hypo-methylated regions in aged sperm

To examine biological significance of the DMRs in aged sperm, we further conducted bioinformatics analyses. Motif analysis revealed significant enrichment (P=1e-25) of a unique consensus sequence, GGAGCTGTCCATGGTGCTGA, which indicates potential binding to the RE1-silencing transcription factor, REST, or in other name, neuron-restrictive silencer factor (NRSF) (Fig. 4d). It should be noted that REST/NRSF is a pivotal regulator for brain development ^16, 17^. Strikingly, these REST/NRSF-binding motifs were detected specifically in the hypo-methylated genomic regions at either of promoter or intron regions (19/96), but not in the hyper-methylated regions (0/16), of aged sperm. As an alternative approach, we used ChIP-Atlas (http://chip-atlas.org), an online database integrating published ChIP-seq data ^43^, to survey transcription factors with enriched binding for hypo-methylated DMRs. Surprisingly, the most significant one was the ChIP-seq data of REST in ES-cell derived neural progenitor cells ^44^; these REST-bound regions were detected in 37/96 of the hypo-methylated genomic regions (Fig.S5, Table S3).

By comparing these 19 motifs and 37 sequences, 15 were common in both analyses (shown in gray in Fig. S5). Interestingly, some of the 37 REST-bound hypo-methylated genomic regions were found to be close to genes associated with autism (SFARI genes) and/or schizophrenia (SZ gene) including *Shank2* and *Htr6*. We further performed gene ontology (GO) analyses using genes within or neighboring to the DMRs. It is revealed that hypo-methylated genomic regions in aged sperm were highly and significantly enriched with genes related to “cell projection”, “learning”, “learning or memory”, “neuron part”, and “connexin complex” (Fig. 4e, Fig. S6a). In MGI’s GO analysis, again, gene lists such as “abnormal nervous system physiology”, “nervous system phenotype”, and “abnormal long term depression” were found in genes related to the hypo-methylated genomic regions (Fig. 4f, Fig. S6b). These results suggest that hypo-methylated genomic regions in aged sperm were enriched with REST/NRSF binding sites and might be related with the neuronal structure and/or function.

### Up-regulation of genes related with ASD-risk and late neurogenesis in the embryonic brain of offspring derived from aged father

Since we found the hypo-methylated genomic regions with REST/NRSF binding motif in aged sperm as well as impaired brain structures in the offspring derived from aged father, we further analyzed gene expression profiles in the developing forebrain of offspring at E14.5 when neuronal production reaches its peak. A volcano plot of RNA-seq data indicates global changes in gene expression in the offspring’s forebrain derived from aged father, but with few significantly up- or down-regulated genes by fold changes of >2.0 or <0.5 with significance less than *p*<0.01 (Fig. 5a). Therefore, we assumed that the cumulative or synergistic effects of small changes in a large number of genes might contribute to the alteration in the brain structure, neuronal activity, and behavior phenotypes.

**Figure 5.**
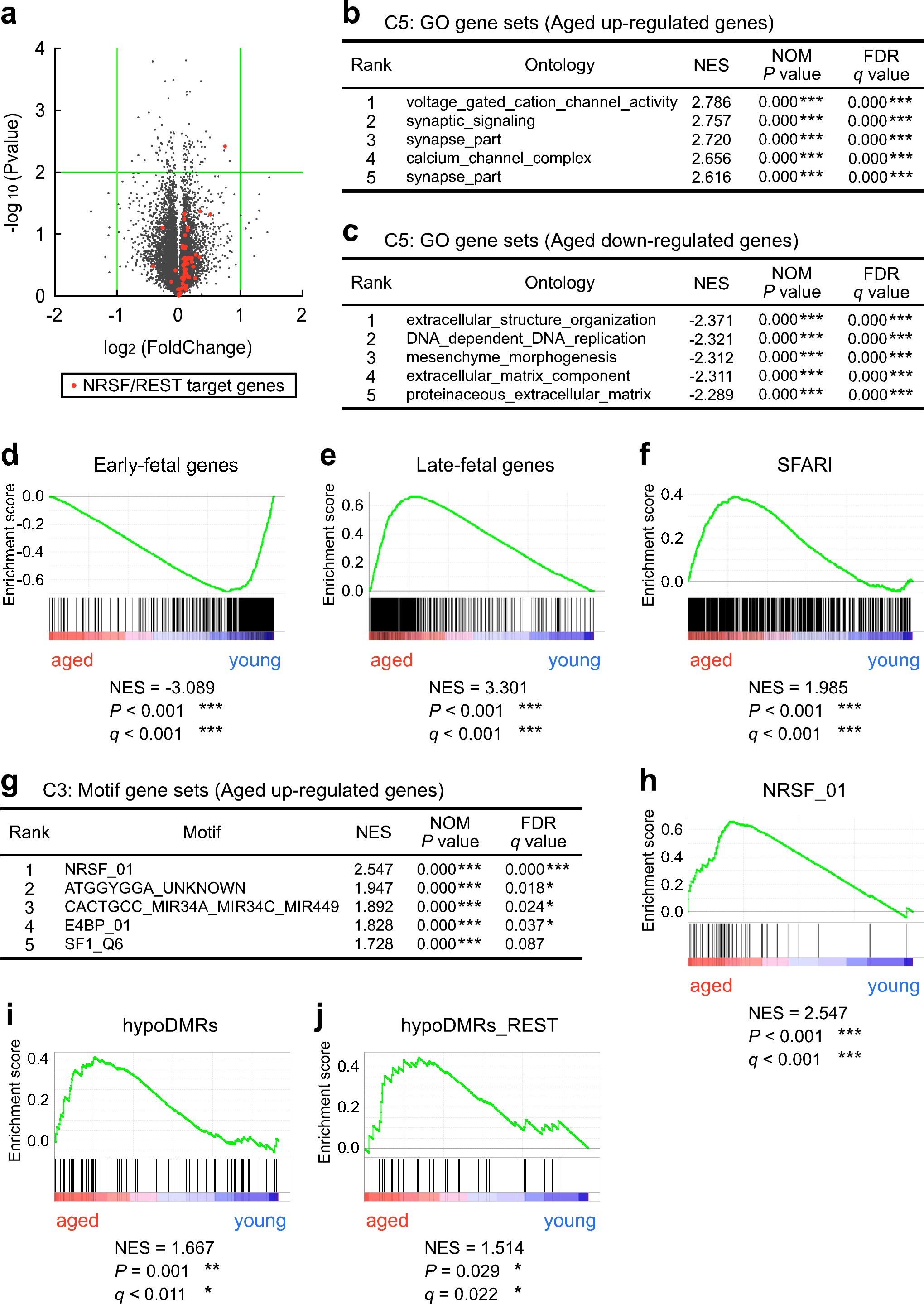
Up-regulation of genes related with late neurogenesis as well as ASD in the developing brain of offspring derived from aged father. (a) Volcano plot for differentially expressed genes in offspring brain derived from aged father at E14.5. Green lines indicate a twofold change and *p*=0.01. Orange dots indicate REST/NRSF target genes. (b, c) GSEA analysis showing GO terms enriched for genes up- or down-regulated in the offspring brain derived from aged father. (d, e) GSEA plots of differentially expressed genes for neural development-related gene sets. NES, normalized enrichment score. (f) GSEA plots of differentially expressed genes for the list of ASD-related gene database, SFARI. (g, h) GSEA analysis showing sequence motifs enriched for genes up-regulated in the offspring brain derived from aged father (g), with the top ranked GSEA plot depicted in (h). (i) GSEA plots of differentially expressed genes for the list of near hypo-methylated genomic regions. (j) GSEA plots of differentially expressed genes for the list of near hypo-methylated genomic regions with REST/NRSF binding motifs. \**p*<0.05, \*\*\**p*<0.001, determined by nominal P value by GSEA.

We next applied Gene Set Enrichment Analysis (GSEA) that is suitable for detecting global changes in gene expression ^45^, and revealed significant enrichment of gene set having GO terms related with neuronal differentiation/function (e.g., voltage gated cation channel activity, synaptic signaling, synapse part, calcium channel complex, synapse, etc) in genes up-regulated in the offspring’s forebrain derived from aged father (Fig. 5b). In contrast, gene set having GO terms related with extracellular structure, DNA replication, and extracellular matrix were listed among the top 5 GO terms was down-regulated in the offspring’s forebrain derived from aged father (Fig. 5c).

Interestingly, we noticed a significant enrichment of the early-fetal gene set ^46, 47^ in the up-regulated genes of the embryonic brain derived from young father (NES=−3.089, *p*<0.001, Fig. 5d). Conversely, a significant enrichment of the late-fetal gene set (NES=3.301, *p*<0.001, Fig. 5e) ^46, 47^ was observed in the up-regulated genes of offspring derived from aged father. It is thus reasonable to assume that precocious neurogenesis occurs in the developing brain derived from aged father. Furthermore, SFARI gene set was also enriched in the embryonic brain derived from aged father (NES=1.985, *p*<0.001, Fig. 5f). It is of note that more SFARI genes (blue dots in Fig. S7a) are plotted in the genes up-regulated in the offspring’s brain derived from aged father, while schizophrenia-related genes (http://www.szgene.org) are evenly plotted in up- and down-regulated genes (Fig. S7b). It should be noted that REST/NRSF target gene set was also enriched in the up-regulated genes in the offspring derived from aged father (Fig. 5g, h, plotted as red dots in Fig. 5a). These suggest that paternal aging can indeed disturb harmonized expression of multiple genes during brain development.

### Up-regulation of genes within/neighboring the hypo-methylated DMRs in the embryonic brain of offspring derived from aged father

To examine impact of the altered DNA methylation in aged sperm on gene expression in the embryonic brain derived from aged father, we listed 94 genes in the vicinity of the hypo-methylated genomic regions (Table S4), and performed quantitative PCR analyses using the F1 forebrain derived from young and aged father at E14.5. As expected, we found much more up-regulated genes than down-regulated ones in the brain of offspring derived from aged father (Fig. S8). Although there were only a few significantly up- and down-regulated genes by fold changes of >2 or <0.5 with significance less than *p*<0.01, the up-regulated genes included ASD and/or schizophrenia-related genes and those with REST/NRSF motif (Fig. S7. Consistently, GSEA using E14.5 brain revealed in the up-regulated genes a significant enrichment of the gene set near the hypo-methylated genomic regions (NES=1.667, *p*=0.001, Fig. 5i). Furthermore, a set of genes near the hypo-methylated genomic regions with REST/NRSF binding motif were enriched in the up-regulated genes (NES=1.514, *p*=0.029, Fig.5j). These suggest an intriguing possibility that DNA hypo-methylation in aged sperm may perturb expression of multiple genes in the offspring’s brain, which might affect abnormal brain structure and behavior.

## Discussion

This study demonstrates that paternal aging affected physical and behavioral abnormalities in their offspring (Fig.1b-f). Less vocal communication, reduced PPI and learning abnormality in F1 offspring derived from aged father are considered to be ASD phenotypes in animal models, while impairment of PPI and learning can also be endophenotypes for schizophrenia ^18^ Therefore, we consider our paternal aging model in mice is suitable for further analyze underlying molecular mechanisms in these psychiatric disorders.

In a previous report using Swiss albino mice showed an opposite tendency in maternal separation-induced USV in offspring; those derived from aged father showed more USV calls ^10^. The discrepancy might be due to difference in genetic background, and/or in environmental factors during paternal aging and/or offspring’s breeding. Indeed, we noticed that the litter size can matter the vocalization of pups; offspring of more than 6 in the litter size showed much vocalization (Fig. S9). Thus, we only used offspring with the litter size of 6 or more in our study.

We found that paternal aging caused low body weight at P6 in mice, as reported in human studies ^23, 24, 25^. This phenotype still persisted in F2 offspring derived from aged F0 grandfather even though they were born to young F1 father. Low body weight at birth or at the early postnatal stage can be a risk for many diseases such as cardiovascular diseases and metabolic syndrome in adult, as proposed in Developmental Origin of Health and Disease (DOHaD) theory ^48, 49^. Here, our findings provide a valuable insight for transgenerational effects of paternal aging on various diseases.

*De novo* mutations in sperm have been paid a great amount of attention as an underlying mechanism of neurodevelopmental disorders due to paternal aging ^12, 13, 49^. In this study, we found that impairment in vocal communication, sensorimotor gating, and spatial learning was canceled in F2 offspring derived from young F1 father born to aged F2 grandfather (Fig. S2c-f). This could rather suggest that transgenerational influence of paternal aging on behavioral abnormalities might not always be transmitted to grandchildren. Therefore, we consider that epigenetic mechanisms may rather be involved in underlying influence of paternal aging, even though a possibility of genetic changes according to father’s age cannot be excluded.

Here we identified, from the comprehensive target methylome analyses, unique 16 hyper- and 96 hypo-methylated DMRs across the chromosomes in aged mouse sperm. Our results are consistent with previous studies using aged sperm in human and rodent, in which hypo-methylated regions are dominantly identified ^11, 14^. A current technical limitation in SureSelect is that we could not identify DMRs in sex chromosomes, on which many genes responsible for neurodevelopmental diseases are located. Nevertheless, we revealed enrichment of REST/NRSF binding motif in hypo-methylated, but not hyper-methylated, genomic regions of aged sperm (Fig.4d). The hypo-methylated genomic regions with REST/NRSF binding motif include risk genes for neurodevelopmental disorders such as ASD and schizophrenia (Fig. S5). Therefore, we can conclude that DNA hypo-methylation in sperm due to paternal aging may actually have a biological significance.

Finally, we examined whether altered DNA methylation due to advanced paternal age has impact on gene expression in the offspring brain. Again, we found that both gene sets for late-fetal genes and REST/NRSF target genes were enriched in the offspring’s brain derived from aged father at E14.5 (Fig. 5a, g, h), but not at E11.5 (Fig. S10). Given the fact that REST/NRSF represses the expression of neuronal genes in neural stem cells ^50^, enrichment of late-fetal gene sets in the offspring’s forebrain derived from aged fathers could be attributed to increased expression of REST/NRSF target genes due to the hypo-methylation. Although it looks puzzling why hypo-methylation of the genomic regions is related with leaky expression of the target genes for REST/NRSF that works as a transcriptional repressor, a recent study has shown that REST/NRSF can rather bind to hyper-methylated regions ^51^; that is, aging-induced hypo-methylation at the REST/NRSF binding sites in the sperm genome might be involved in causing de-repression of inadequate gene expression in the developing brain derived from aged father.

A previous study reported reduction of DNA methylation in both aged sperm and the offspring brain from aged father, though no significant relationship was found between DNA methylation and gene expression in the offspring brain ^11^. One possible explanation for the discrepancy for their results and ours is due to a difference of analyses; we adopted a comprehensive GSEA that is suitable for detecting global changes in gene expression ^45^, while they used a conventional correlation analysis of Pearson between DNA methylation and expression of limited genes ^11^. Using this advanced technology, we first uncovered an intriguing possibility that DNA hypo-methylation due to paternal aging indeed has impact on gene expression in the offspring brain.

In another model for ASD using *Chd8*+/ΔL mutant, GSEA has been applied because expression of genes were almost unchanged in the developing brain ^47^. In their results, expression of REST/NRSF is reduced, resulting in increase and reduction of early- and middle-fetal genes, respectively ^47^. We did not find change in REST/NRSF expression in our RNA-seq data using developing brains neither at E11.5 nor E14.5 (Fig. S11). Phenotype of the brain structure also seem to be opposite; *Chd8*+/ΔL mutant showed a bigger brain, while offspring derived from aged father exhibited a reduction of cortical thickness. Therefore, although there may be different regulatory mechanisms, both up- or down-regulation of REST/NRSF target genes during brain development might be a common risk for neurodevelopmental disorders.

To summarize, we would like to propose a model that paternal aging may induce leaky expression of REST/NRSF target genes that have been marked with hypo-methylation within a sperm cell, causing precocious neurogenesis, thereby resulting in abnormality in brain structures and neuronal activities, which may cause behavior phenotypes reflecting some of the phenotypes of neurodevelopmental disorders (Fig.6). The advantage of this scenario is that it can explain individual differences and multigenetic contributions in comorbid symptoms of individual patients. It will be a next challenge to elucidate whether DNA hypo-methylation of REST/NRSF binding motif in sperm due to paternal aging impacts expression of REST/NRSF target genes in embryonic brain at the individual level and to reveal whether hypo-methylation of the sperm DNA might induce abnormalities in offspring’s behavior. Although there still is a mystery how de-methylated marks in the sperm genome can influence gene expression in the offspring brain, a recent etiological data suggest a possible involvement of methylation levels in regard with the risk for ASD ^52^.

**Figure 6.**
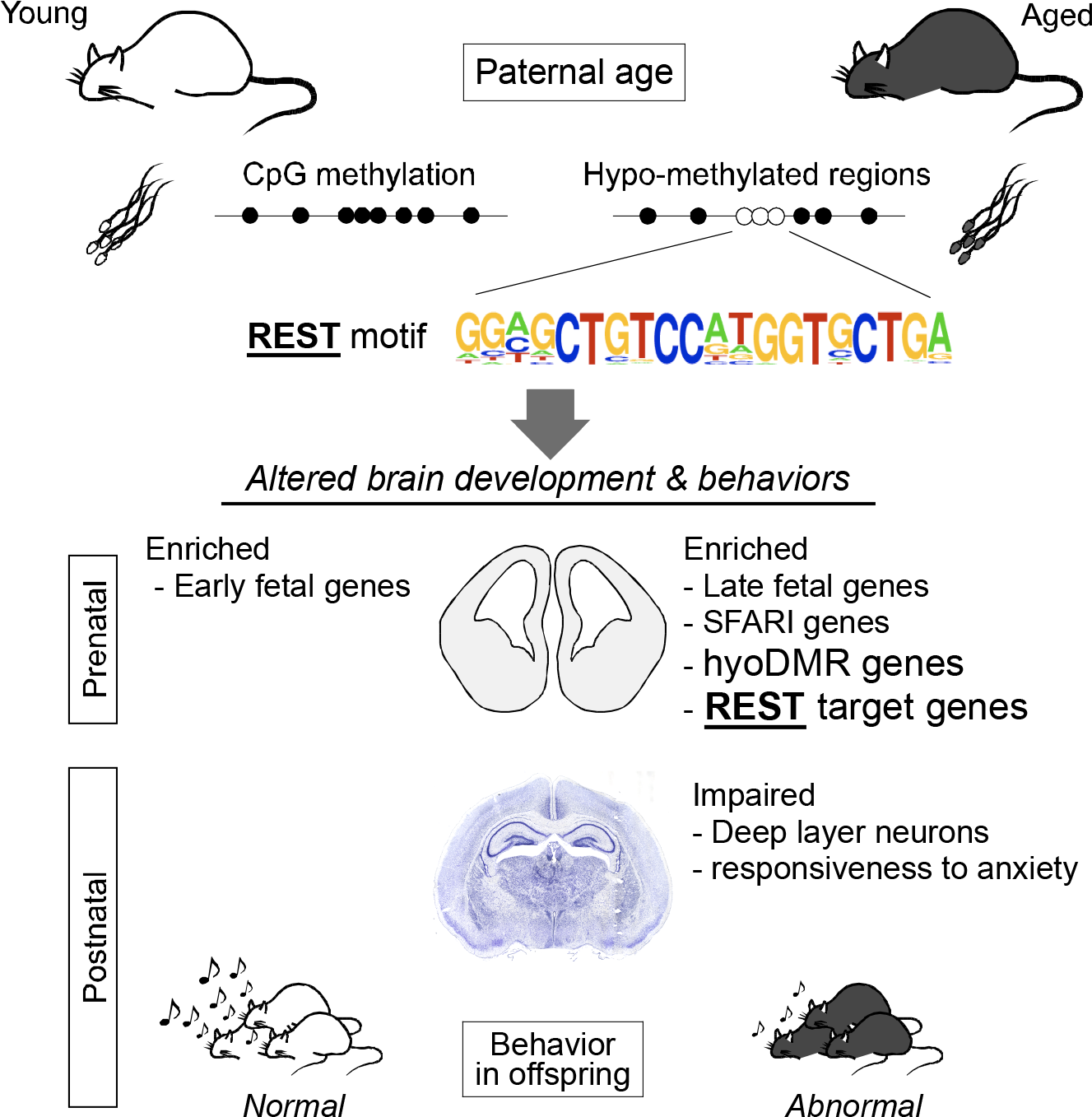
Graphical summary. Possible mechanism of neurodevelopmental and behavioral abnormalities due to paternal aging. Paternal age induces in the sperm genome hypo-methylation especially in regions with REST/NRSF binding motifs in sperm, leading to leaky expression of the REST/NRSF target genes in the developing brain of the offspring, eventually causing abnormalities in the brain structure and neuronal activity. Potential variety of each DMR may reflect diversity in neurodevelopmental disorders at the individual level.

## Materials and Methods

### Animals

All experimental procedures were approved by Ethic Committee for Animal Experiments of Tohoku University Graduate School of Medicine (#2014-112) as well as of Tokyo University of Agriculture, and animals were treated according to the National Institutes of Health guidance for the care and use of laboratory animals. Three-month-old (young) or >12-month-old (advanced-age) male C57BL6/J mice were crossed with young virgin female mice to obtain F1 offspring for this study (Fig.1a). To obtain F2 offspring, young F1 father derived from young or aged grandfather were mated with young virgin female mice (Fig.2a). On postnatal day 6, some offspring were tattooed with an Aramis Animal Microtattoo System (Natsume Co., Ltd., Tokyo, Japan) for individual identification in behavior analyses in adult. Male mice were separated from female mice 1 week after mating to minimize confounding factors for behavior of the offspring. All animals were housed in standard cages in a temperature and humidity-controlled room with a 12-h light/dark cycle (light on at 8:00) and had free access to standard lab chow and tap water. For DNA methylation analyses, sperm was obtained from 3-month or 18-month-old male mice. The number of mice used for comprehensive behavior analyses and qPCR is indicated in Supplemental Table 1.

### Behavior tests

#### Ultrasonic vocalization (USV)

Each pup was assessed for USV on postnatal day 6 (P6) according to previously described protocols ^19, 20, 21, 22^. The pup was separated from its mother and littermates, one at a time, and placed in a plastic dish with floor cloth in a soundproof chamber, and USV calls were recorded for 5 min with a microphone connected with the UltraSound Gate 416H detector set (Avisoft Bioacoustics, Germany) at 25-125 kHz to measure the number of USV calls and latency until first USV calls. Since we found that the litter size influences on pups’ USV calls, we only used data of the offspring whose litter size is 6 and above (Fig.S9).

#### Prepulse inhibition (PPI) test

PPI test was conducted as described previously ^53^. The mice were introduced into a plastic cylinder of a startle chamber (SR-LAB, San Diego Instruments, San Diego, CA). After 5 min acclimatization with 65 dB broadband background noise, PPI test session with total of 64 trials was conducted. The test session was consist of five types of trials as follows: no stimulus trials (background noise only); startle pulse alone trials (40 ms duration at 120 dB, p120); and three prepulse+pulse trials (20 ms duration prepulse at 68 dB, pp3; 71dB, pp6; or 77dB, pp12, followed by a 40 ms duration startle stimulus at 120 dB after a 100 ms delay). Each trial was presented in a pseudo random manner by SR-LAB software system. The startle response was detected by an accelerometer under the plastic cylinder and recorded as electronic signals.

#### Morris water maze test

Morris water maze test was performed as described previously with some modifications ^54^. Briefly, circular pool with 1 m diameter was filled with opaque water and the water temperature was maintained at 22°C using a thermostat. The pool was surrounded by curtain screens that had distinctive markings on the surface to enhance spatial location. The pool was monitored by a video camera connected to auto tracking system (WaterMaze, ActiMetrics, IL). In this study, we trained mice for 9 days to test spatial learning and reversal learning. First, the mice were trained for 4 days in the pool to locate a transparent circular platform with 10 cm diameter that was submerged 1-1.5 cm beneath the surface of the water. During the training period, the mice experienced two trials of the swimming session per day. On the day 5, the mice freely swam in the pool without the platform for 1min to test their spatial memory. From day 6, the mice were trained for 3 days in the pool to locate a new position of the platform that was moved to opposite side from the first position. On the day 9, the mice freely swam in the pool without the platform for 1 min to test their ability to reverse a spatial memory once established.

#### Courtship USV

Courtship USV recording was performed with previous report with some modifications ^55^. Female mouse in their own home cage was placed in a recording box for 60 sec. Subsequently, an unfamiliar intruder male mouse was placed into the recording box with resident female mouse, and the ultrasonic vocalization (USV) was recorded for 60 sec. from first USV calls using the UltraSound Gate 416H detector set (Avisoft Bioacoustics, Germany) at 25-125 kHz.

#### Three-chamber social interaction

Three-chamber social interaction test was performed with according to previously report ^56^. Each chamber was 40 × 60 × 20 cm, and the dividing into three small compartments with clear Plexiglas wall, with small square openings (7 × 7 cm) allowing access into each chamber. An unfamiliar C57BL6/J male (stranger) that had no prior contact with the subject mice was placed in one of the side chambers. The location of stranger in the left versus right side chamber was systematically alternated between trials. The stranger mouse was enclosed in a small, round cage, which allowed olfactory, visual, auditory and tactile contacts but did not allow deep contacts. The subject mouse was first placed in the center chamber for 5 min. and then allowed to explore the entire social test apparatus without any stranger mouse for 10 min. Then the mice were explored the apparatus with stranger mouse in a side chamber. Time spent in each chamber was recorded using Any-maze (Brain Science. Idea. Co., Ltd., Japan).

### Target methylome analysis

#### DNA preparation

Sperm DNA was isolated by a standard phenol-chloroform extraction procedure with dithiothreitol. One microgram of DNA was dissolved in 130 μl of 10 mM Tris–HCl (pH 8.0) and sheared with an S220 focused ultrasonicator (Covaris, Woburn, MA, USA) to yield 500-bp fragments. The AMPure XP system (Agilent Technologies, Santa Clara, CA, USA) was used to purify the fragmented DNA as follows. Sheared DNA (130 μl) was mixed with 1.8 × volume (234 μl) of AMPure XP reagent and allowed to stand for 15 min at room temperature. Beads were collected using a magnetic stand, supernatant was removed, and the resulting pelleted beads were rinsed with 70% ethanol and dried by incubation at 37°C for 5 min. DNA was then eluted from the beads with 20 μl of RNase-free water. The eluted DNA solution was dried under vacuum and dissolved in 7 μl of RNase-free water.

#### Target enrichment

Target enrichment by liquid-phase hybridization capture was performed using the SureSelect Mouse Methyl-Seq kit (Agilent Technologies) ^42^. Genomic DNA (7 μl) that was fragmented and purified as described above was supplemented with 3 μl of formamide (biochemistry grade; Wako Pure Chemical Industries, Osaka, Japan) and overlaid with 80 μl of mineral oil (Sigma-Aldrich, St. Louis, MO, USA). DNA was completely denatured by incubating the tube at 99°C for 10 min, then the tube was cooled to 65°C and maintained at that temperature for at least 5 min before adding the following reagents. Hybridization buffer and capture probe mix were prepared according to the manufacturer’s protocol, and were each overlaid with 80 μl of mineral oil and incubated at 65°C for 10 min. The two solutions were then combined and mixed thoroughly by pipetting. The combined solution was transferred to a tube containing the denatured input DNA (maintained at 65°C as described above) and the solution was thoroughly mixed by pipetting. The tube was incubated at 65°C for 24 h to allow hybridization between probes and targets. A 50 μl volume of well-suspended DynaBeads MyOne Streptavidin T1 solution (Life Technologies, Carlsbad, CA, USA) was placed in a 1.5 mL tube, and the beads were washed twice with 200 μl of binding buffer. The hybridization reaction, supplemented with 200 μl of binding buffer, was added to the pelleted beads and thoroughly mixed. After incubation at room temperature for 30 min with agitation, the beads were collected using a magnetic stand and washed with 500 μl of wash buffer 1, and then subjected to three rounds of washing and re-suspension in pre-warmed buffer 2, followed by incubation at 65°C for 10 min. After removing the washing solution from the tube, enriched DNA was eluted by incubating the beads in 20 μl of elution buffer at room temperature for 20 min. The eluate was immediately subjected to bisulfite treatment.

#### Bisulfite treatment

The EZ DNA Methylation-Gold kit (Zymo Research, Irvine, CA, USA) was used for bisulfite treatment of target-enriched DNA according to the manufacturer’s instructions. The enriched DNA solution (20 μl) was mixed with 130 μl of freshly prepared CT conversion reagent, and the mixture was incubated at 64°C for 2.5 h. The 10 min incubation step at 98°C was omitted, since the target-enriched DNA was already denatured. After purification and desulfonation, bisulfite-treated DNA was eluted with 20 μl of M-elution buffer.

#### PBAT library construction and Illumina sequencing

We used bisulfite-treated DNA for library preparation according to the PBAT protocol^24^ (also available from http://crest-ihec.jp/english/epigenome/index.html), except that the following primers were used. The primer used for first-strand synthesis was 5′-biotin ACA CTC TTT CCC TAC ACG ACG CTC TTC CGA TCT WWW WNN NN-3′ (W = A or T). The indexed primer used for second-strand synthesis was 5′-CAA GCA GAA GAC GGC ATA CGA GAT XXX XXX GTA AAA CGA CGG CCA GCA GGA AAC AGC TAT GAC WWW WNN NN-3′, where XXX XXX represents the index sequence of each primer. Constructed SSM-PBAT libraries were sequenced as previously described ^39, 41, 57^ using the Illumina HiSeq2500 system (San Diego, CA, USA) to generate 100-nt single-end sequence reads. Before alignment, each random sequence are trimmed from the sequence data sets. These trimmed data (93-nt) are available with the DNA Data Bank of Japan (DDBJ) accession number DRA007933.

#### Target methylome sequence alignment and statistical analysis

SSM-PBAT reads were aligned to the mouse genome (mm10, Genome Reference Consortium Mouse Build 38) using the Bismark tool (v.0.10.0; http://www.bioinformatics.babraham.ac.uk/projects/bismark/) with the specific options: −q −n 2 ‒l 93 ‒pbat. Statistical significance of DNA methylation at each CpG site and CGI was evaluated by Model-based Analysis of Bisalfite Sequence data (MOABS). Composite profiles were generated using SeqMonk (http://www.bioinformatics.babraham.ac.uk/projects/seqmonk/).

### Immunohistochemistry

Procedures for immunohistochemistry were according to previous studies ^58, 59^. Coronal serial sections (16 mm thick) were cut on a cryostat, washed with Tris-based saline containing 0.1% tween20 (TBST) and treated with 3% bovine serum albumin and 0.3% TritonX-100 in PBS solution (blocking solution). The sections were incubated with primary antibodies against Tbr1 (AB10554; Millipore, Morcocco; diluted in 1: 200), Satb2 (ab51502; Abcam, Cambridge, UK; diluted in 1:1,000), Ctip2 (ab18465; Abcam, Cambridge, UK; diluted in 1:1,000), Cux1 (sc-13024; Santacruz Biotechnology, Dallas, US; diluted in 1:500) in blocking solution overnight at 4°C. Subsequently, the sections were incubated with Cy3 or Alexa488 conjugated secondary antibodies (Jackson ImmunoResearch Inc.; diluted 1:500) and 4’, 6-diamino-2-phenylindole (DAPI; diluted in 1:1,000) for 1 hour at room temperature. Images were obtained by using a fluorescent microscopic system (BZ-X, Keyence, Osaka, Japan). Cortical thickness was measured by an analyzing tool implemented on the system.

### Gene expression analyses

#### RNA extraction for RNA sequencing

Total RNA was extracted from embryonic telencephalons at E11.5 and E14.5 using RNeasy Micro (Qiagen, Germany) according to the manufacturer’s protocol. The libraries were obtained by 2.0 μg of total RNA using TruSeq^®^ Stranded mRNA Library Prep Kit (Illumina, USA) following the guideline of the kit. The quality of libraries was determined by an Agilent 2100 Bioanalyzer (Agilent Technologies, Germany). Cluster generation of the library was performed on a cBot with Illumina HiSeq PE Cluster Kit v4 – cBot – HS (Illumina, USA). The libraries were sequenced following a paired-end method (100 base pair reads) using HiSeq 2500 and HiSeq Control Software v2.2.58/Real Time Analysis v1.18.64 (Illumina, USA). BCL files were converted to FASTQ file formats by using bcl2fastq2 (Illumina, USA).

#### Analysis of RNA sequencing data

A series of RNA-seq data analyses were performed by the instruction of Genedata Profiler Genome (Genedata). The reads in FASTQ format were aligned to the reference of mouse genome (GRCm38/mm10, Dec.2011) with TopHat v2.0.14, Bowtie, and SAM tool ^60, 61, 62, 63^. These filtered data are available with DNA Data Bank of Japan (DDBJ) accession number DRA008023. Unreliable filtered data was removed: the dispersion of fragments per kilobase of transcript per Million mapped reads (FPKM) was zero and the number of reads was no more than 16 in samples.

#### Enrichment analysis

Genome coordinates of DMRs and other SureSelect target regions were converted from mm10 to mm9 assembly with UCSC liftOver tool (https://genome.ucsc.edu/cgi-bin/hgLiftOver), prior to enrichment analyses with ChIP-Atlas. Enrichment analyses of GO terms (http://geneontology.org) and MGI phenotypes (http://www.informatics.jax.org) were performed for genes locating nearby DMRs against the other RefSeq coding genes. GSEA was performed according to Katayama et al 2016, and SFARI gene set ^64^ and SZ gene sets ^65^ were obtained from https://www.sfari.org and http://www.szgene.org, respectively.

### Statistic analyses

All data are presented as a dot plot or mean ± standard error of the mean (SEM). Statistical analysis was performed using Student *t* test, Wilcoxon, two-way ANOVA, or Tukey-Kramer test. SSPS 16.0 software (SPSS Inc., Chicago, USA) was used for statistic analyses and a *p* value less than 0.05 was considered to be statistically significant.

## Data Availability

DNA methylome data in young and aged sperm that support the findings of this study have been deposited in DNA Data Bank of Japan (DDBJ) with the accession number “DRA007933”.

RNA-sequence data in developing brain derived from young and aged father that support the findings of this study have been deposited in DDBJ with the accession number “DRA008023”.

## Author contribution

### Conception and design of the study

Kaichi Yoshizaki, Ryuichi Kimura, Yasuhisa Matsui, Tomohiro Kono, Noriko Osumi

### Analysis and interpretation of data

Kaichi Yoshizaki, Tasuku Koike, Ryuichi Kimura, Takako Kikkawa, Shinya Oki, Kohei Koike, Kentaro Mochizuki, Hitoshi Inada, Hisato Kobayashi, Yasuhisa Matsui, Tomohiro Kono, Noriko Osumi

### Collection and assembly of data

Kaichi Yoshizaki, Tasuku Koike, Ryuichi Kimura, Takako Kikkawa, Kohei Koike, Kentaro Mochizuki, Hisato Kobayashi

### Drafting of the article

Kaichi Yoshizaki, Ryuichi Kimura, Takako Kikkawa, Shinya Oki, Koike Kohei, Hisato Kobayashi

### Critical revision of the articles for important intellectual content

Yasuhisa Matsui, Tomohiro Kono, Noriko Osumi

### Final approval of the article

Noriko Osumi

### Grant information

16H06530 (JSPS KAKENHI)

P17gm0510017h (AMED-CREST)

---

**Figure S1.**
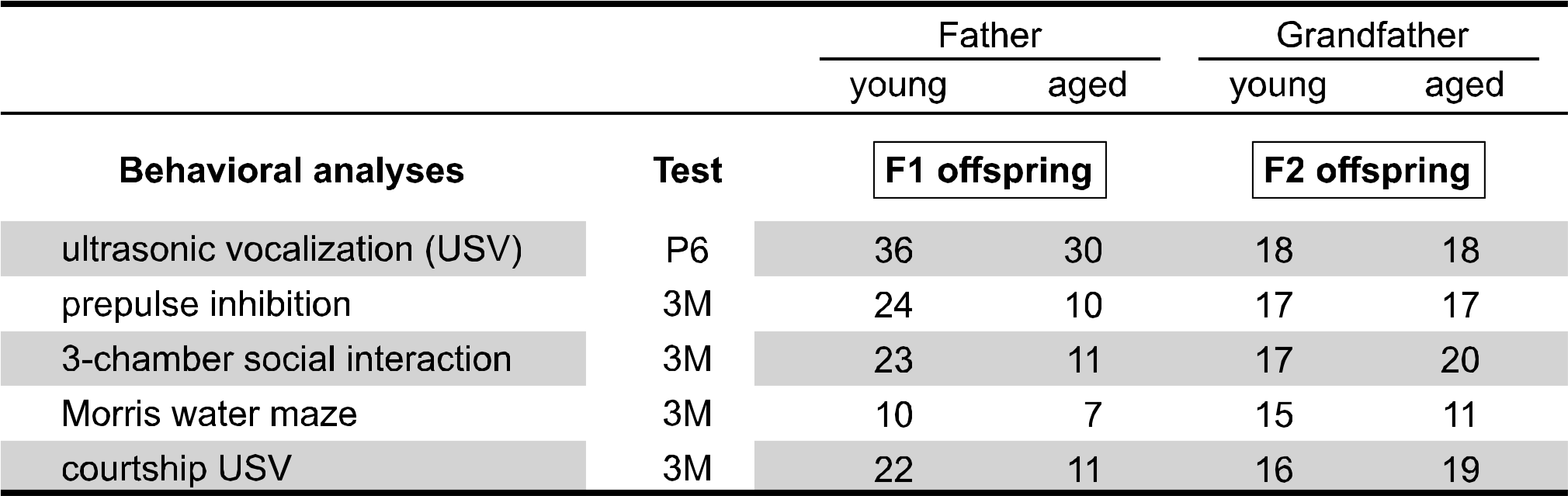
Number of animals used in behavioral analyses. Animal numbers used for each behavioral analysis in F1 offspring (derived from young and aged father) and F2 offspring (derived from young and aged grandfather). Each behavioral analysis was performed at postnatal day 6 or 3-month-old mice.

**Figure S2.**
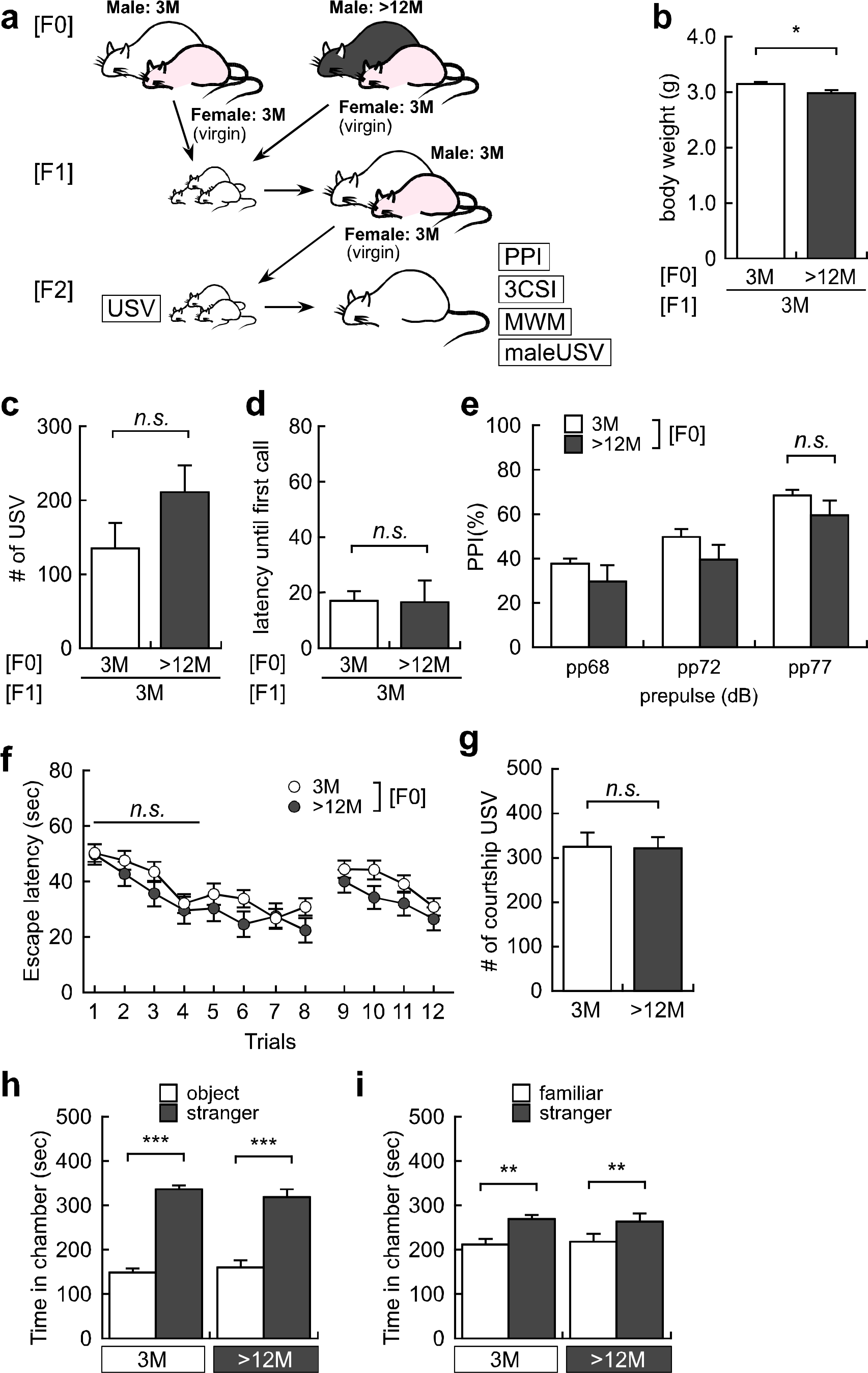
Advanced grand-paternal age affects body weight, but not behavior phenotypes in F2 offspring. (a) Experimental design of F2 offspring derived from young father born to either young or aged grandfather. (b) Body weight, (c) the number of maternal separation-induced ultrasonic vocalization (USV) calls, and (d) latency until first USV calls at postnatal day 6, (e) pre-pulse inhibition score at pp68 dB, pp72 dB and pp77 dB, (f) spatial and reversal learning in Morris water maze test, (g) courtship USV to female mouse, and (h) sociability and (g) social novelty in three chamber social interaction test at 3-month-old in F2 offspring derived from young father born to either young or aged grandfather. All data are presented by the mean ± SEM. \**p*<0.05, \*\**p*<0.01, \*\*\**p*<0.001, determined by Student *t* test or two-way ANOVA followed by post hoc test with Bonferroni method.

**Figure S3.**
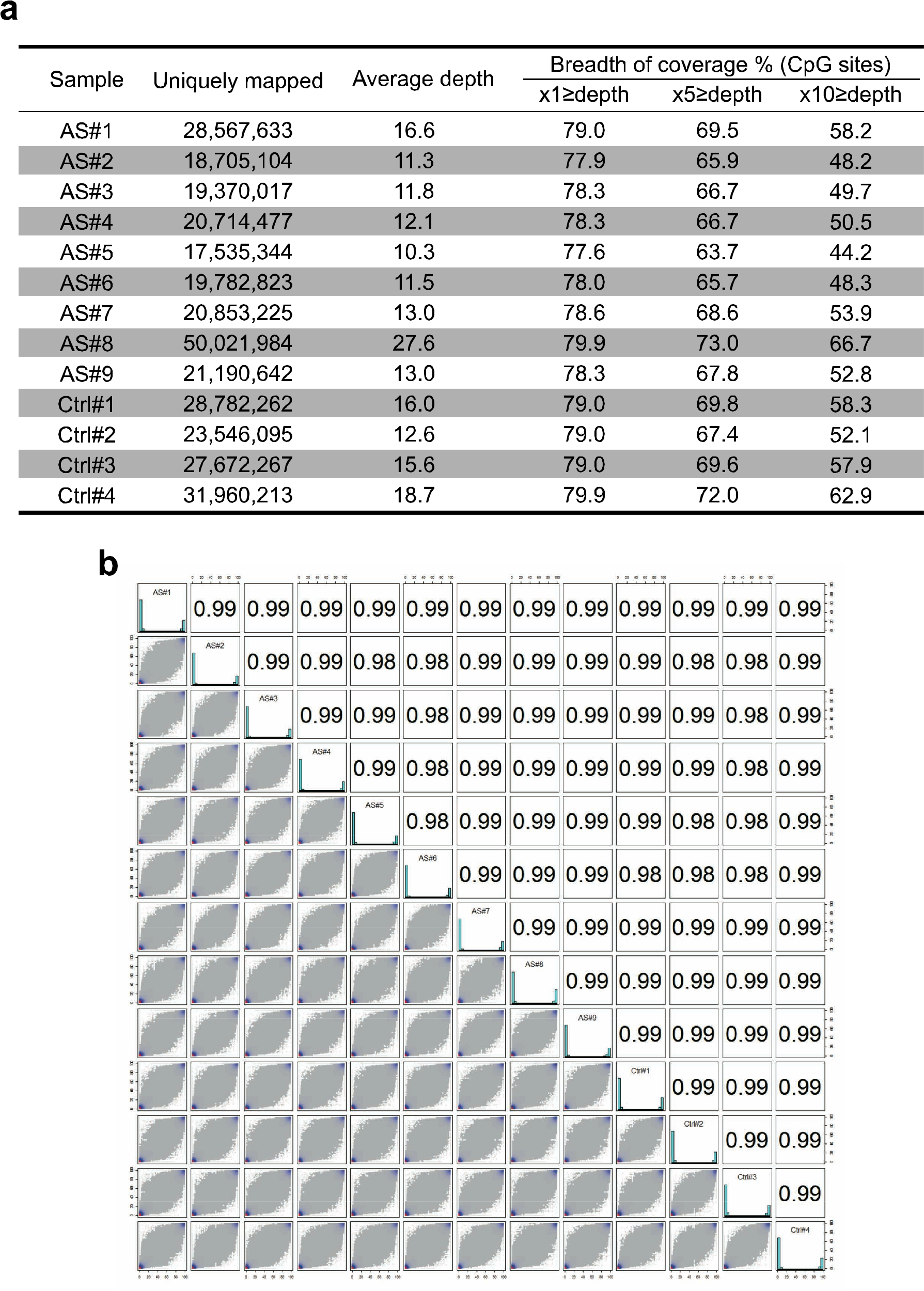
Overall methylation profiling quality is sufficient for further analysis. (a) The uniquely mapped reads, average depth and coverage of each sperm samples for sequence analysis. (b) Correlation coefficients of DNA methylation data.

**Figure S4.**
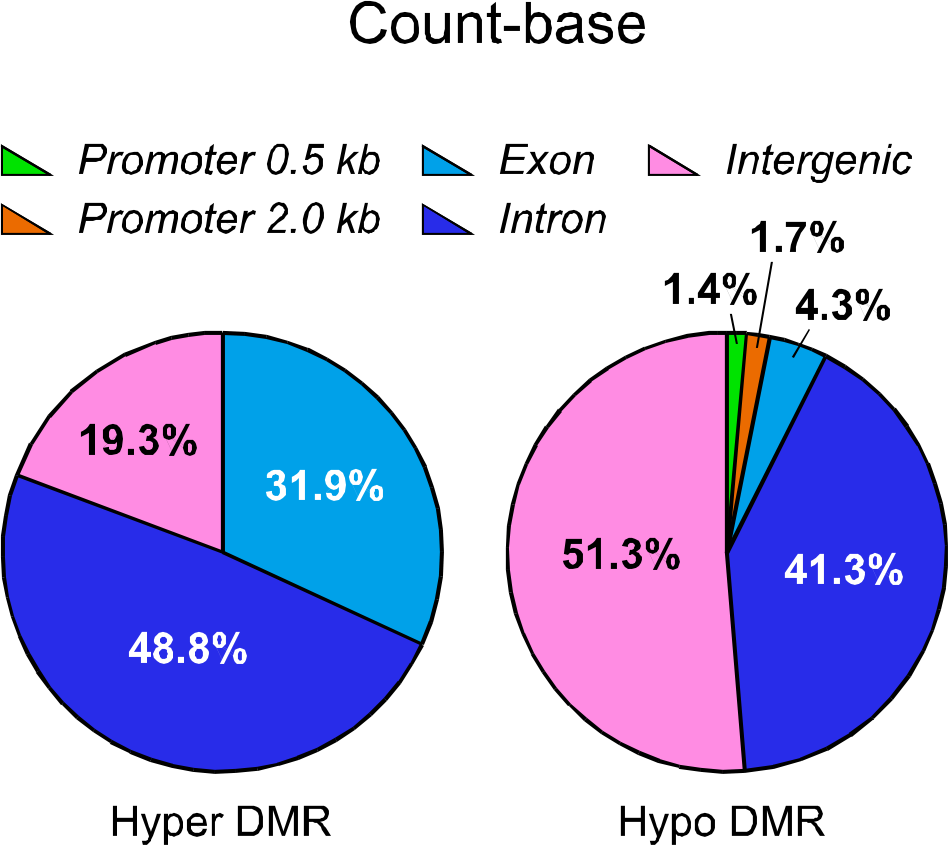
Genomic distribution of age-related hyper and hypo-methylation in sperm. Pie chart showing the distribution of hyper- (left) and hypo- (right) methylated genomic regions (promoter [0.5 kb or 2.0 kb upstream of TSS], exons, introns, intergenic regions) in count-base analyses.

**Figure S5.**
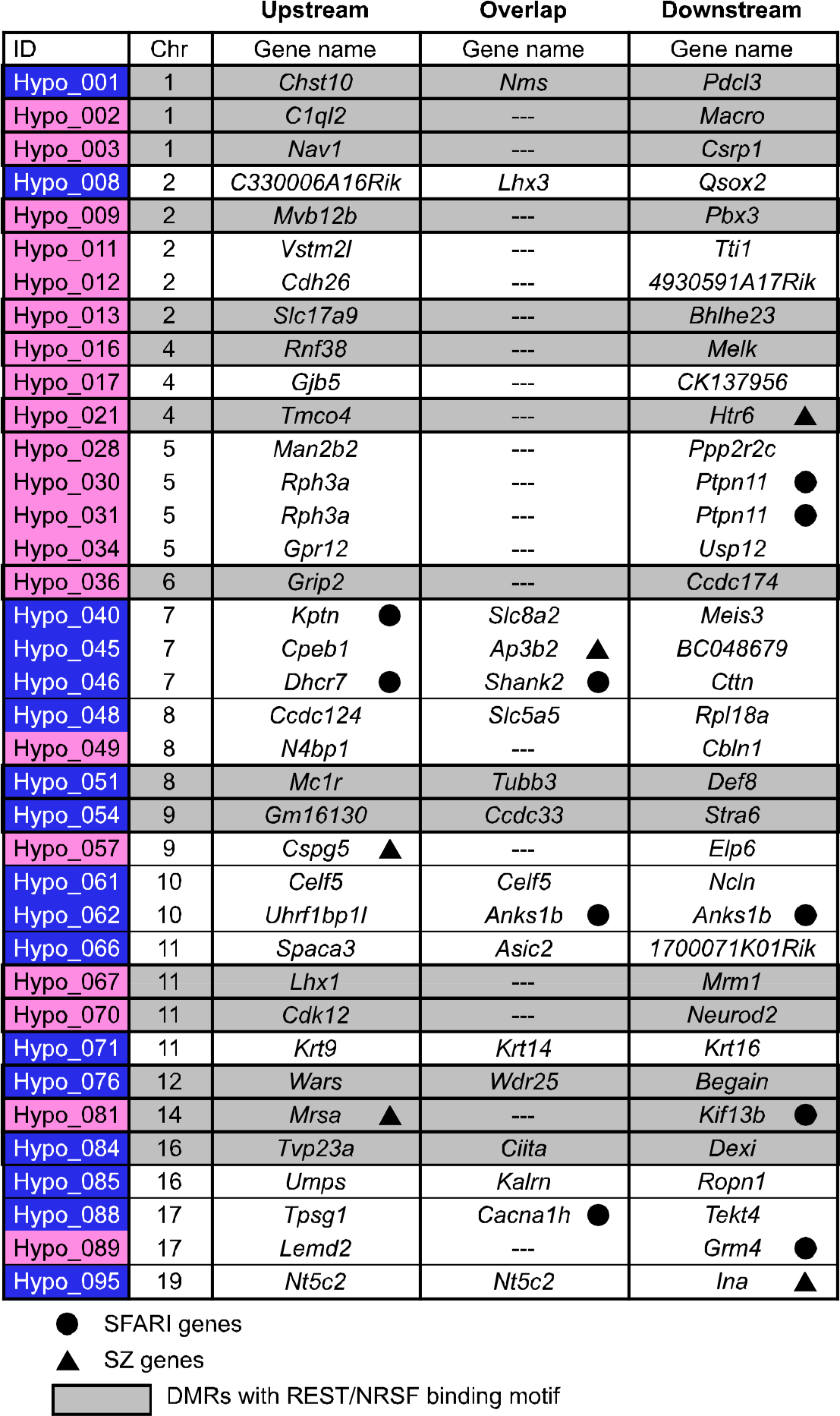
Gene lists near 37 hypo-methylated genomic regions with REST/NRSF binding confirmed by ChIP-Atlas database. Genes near thirty-seven DMRs selected by REST/NRSF binding motifs in embryonic stem cells-derived neural progenitor cells in ChIP-Atlas database were listed from 96 hypo-methylated regions in aged sperm. Blue highlight indicates DMRs in exon of overlap genes. Mazenta highlight indicates DMRs in intergenic regions between upstream and downstream genes. Gray highlight incidates DMRs with REST/NRFS binding motif. Black circle indicates ASD-related genes (SFARI gene) and black triangle indicates schizophrenia-related genes (SZ gene)

**Figure S6.**
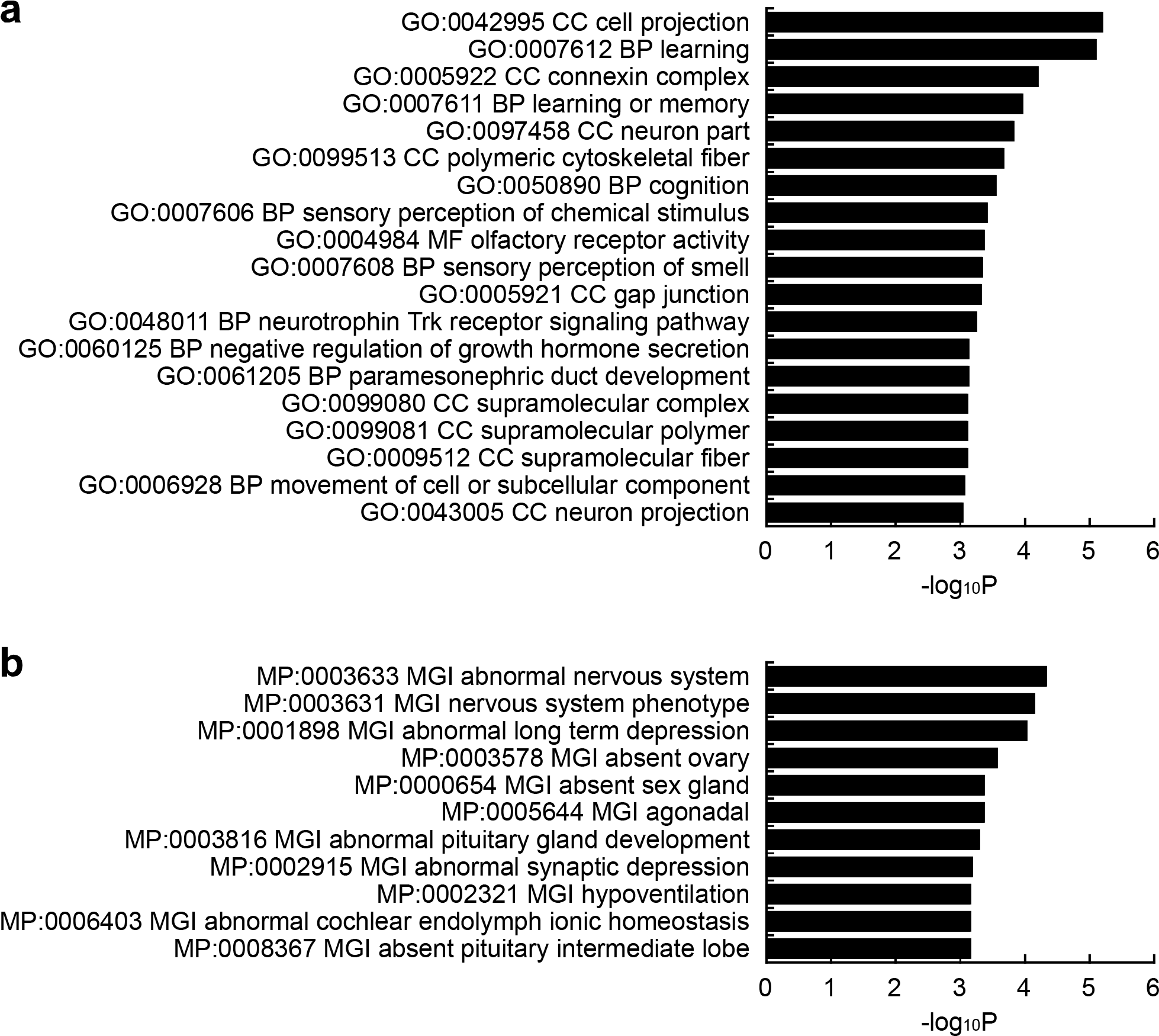
Gene ontology analysis for hypo-methylated DMRs. (a) The top 19 GO terms (*p*<0.001) associated with biological process (BP), cellular component (CC) and molecular function (MF), and (b) the top 11 MGI phenotypes (*p*<0.001) of genes near hypo-methylated DMRs in aged sperm are shown. Both indicate that genes associated with neuronal function are enriched.

**Figure S7.**
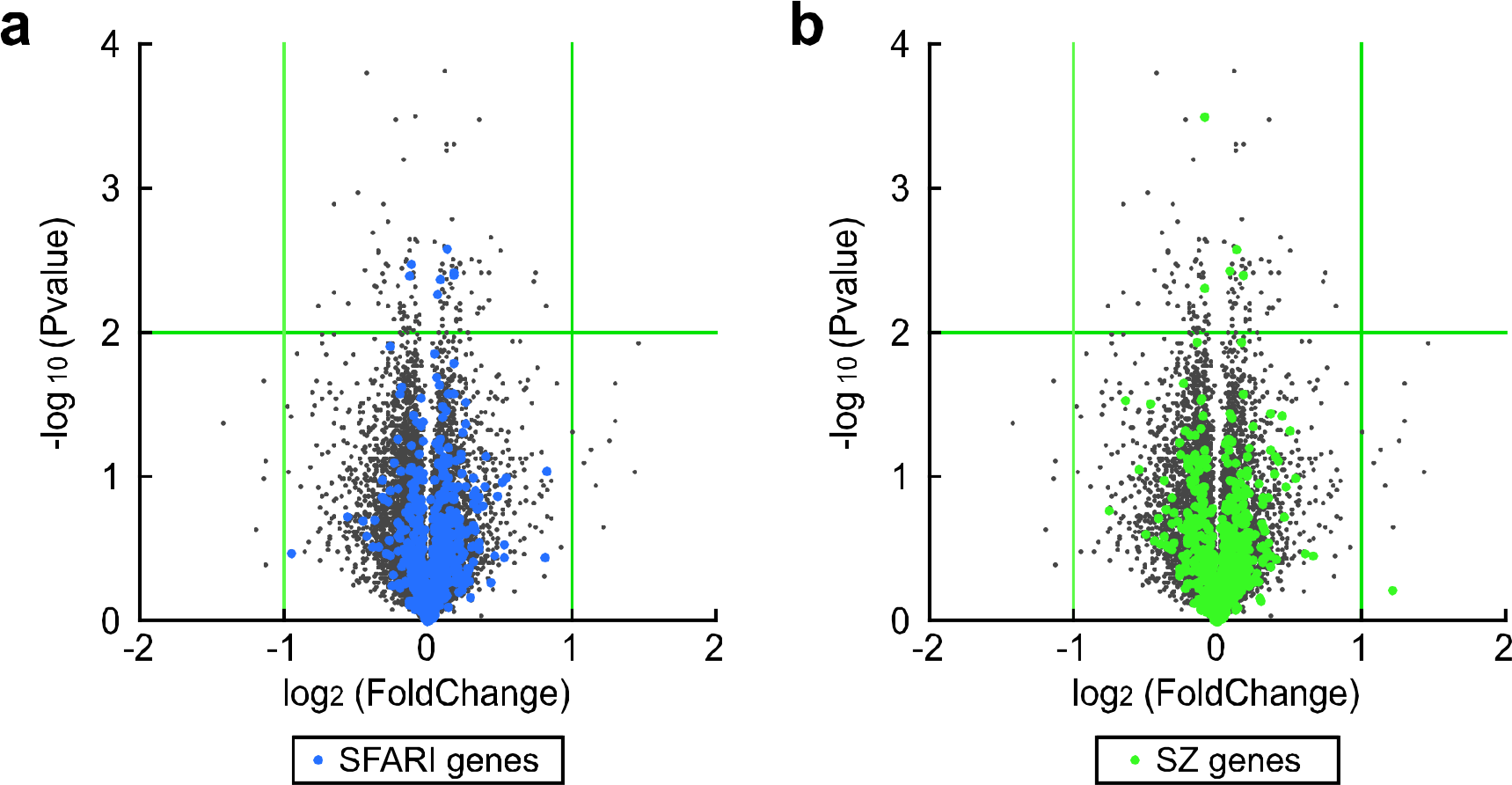
Up- or down-regulation ASD and schizophrenia risk genes in developing brain derived from aged father. (a, b) Volcano plot for differentially expressed genes. Green lines indicate a two-fold change and *p*=0.01. (a) Blue dots indicate ASD-risk genes listed in SFARI database. (b) Green dots indicate schizophrenia-related genes listed in SZ genes database. SFARI genes include more up-regulated ones in expression in the forebrain of E14.5 mouse embryos derived from aged father.

**Figure S8.**
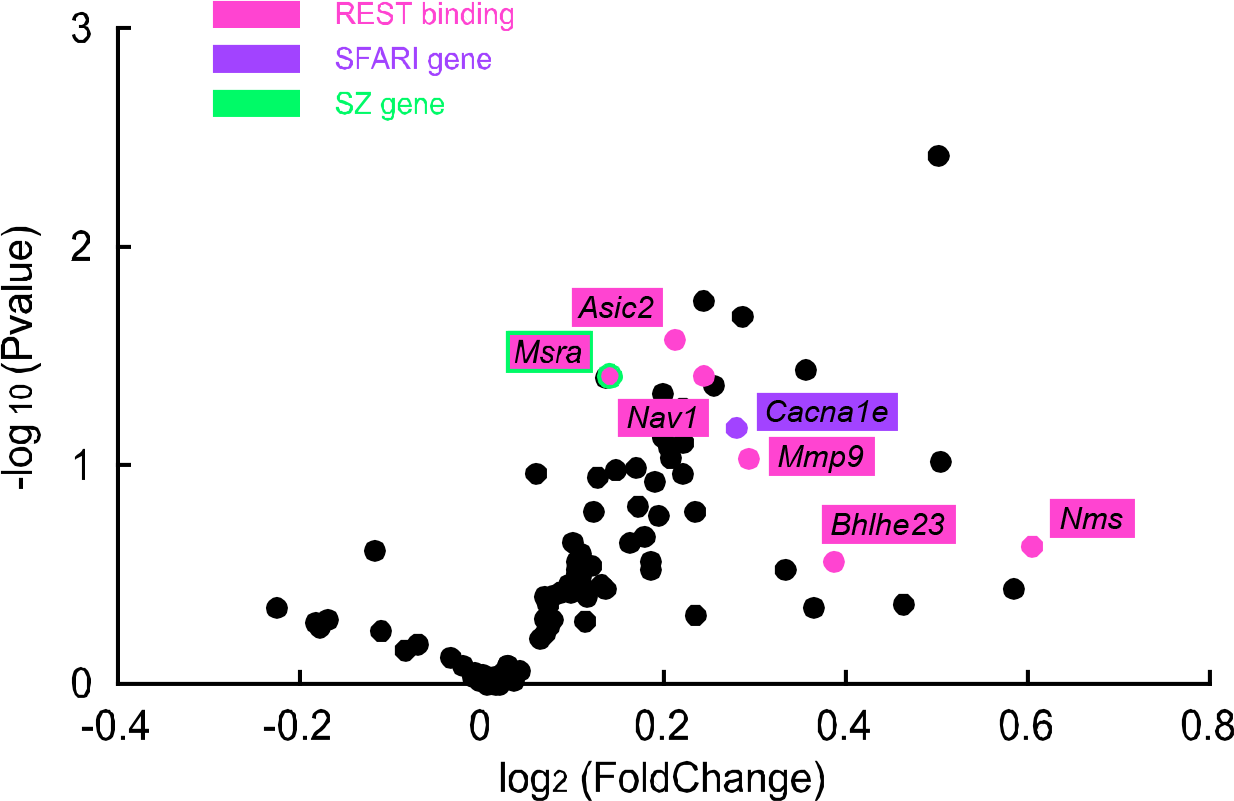
Up-regulation of REST/NRSF target, ASD risk and late neurogenesis genes in the developing brain derived from aged father. (a) Volcano plot for differentially expressed genes in the offspring brain derived from young and aged father at E14.5. Magenta indicates REST binding genes. Purple indicates ASD-related genes (SFARI gene). Green indicates schizophrenia-related genes (SZ gene).

**Figure S9.**
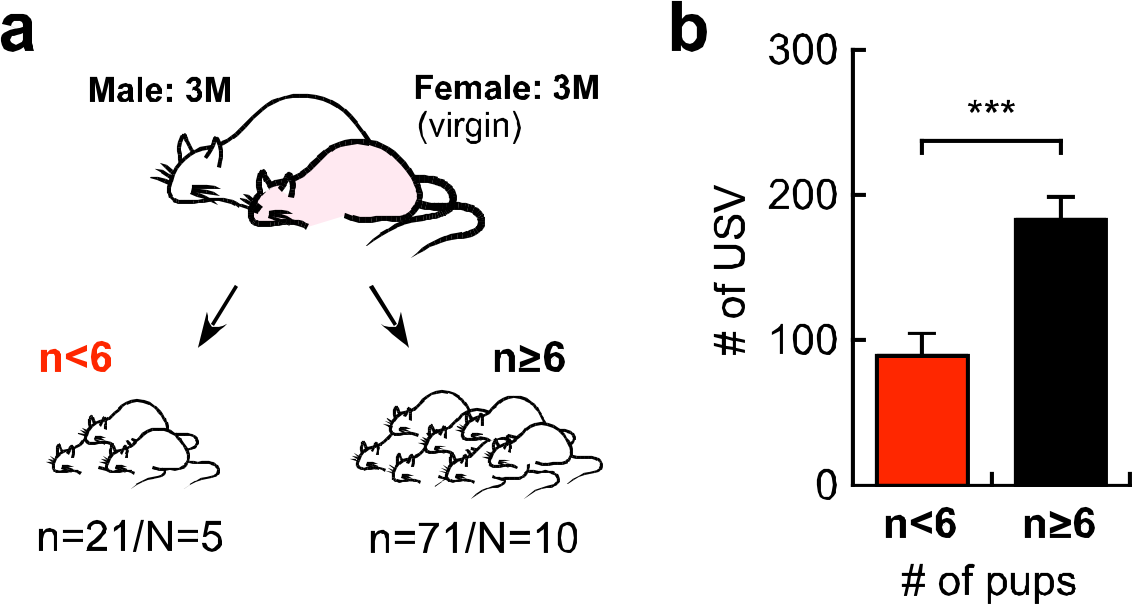
The number of pups affects vocal communication. (a) Experimental design of F1 offspring derived from young father. (b) Number of maternal separation-induced ultrasonic vocalization (USV) calls in offspring in which litter size was 6 and more or 5 and less. Pups with the larger litter size emit more USV calls. Data are presented by the mean ± SEM. \**p*<0.05, determined by Student *t* test.

**Figure S10.**
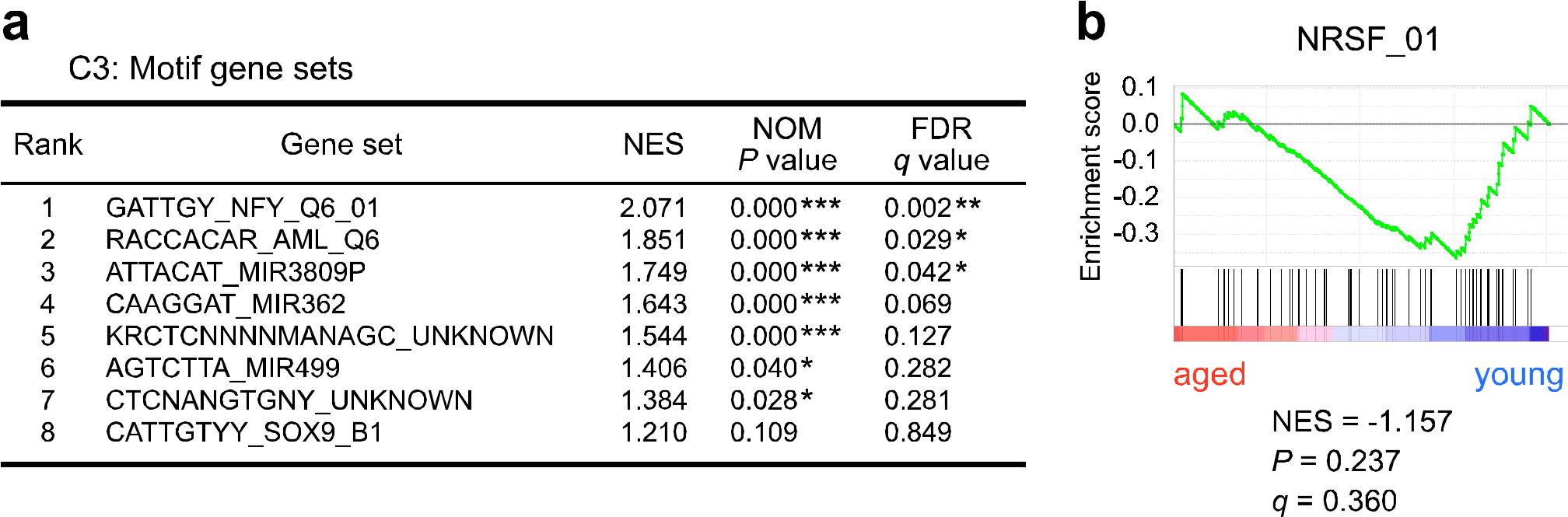
REST/NRSF target genes were not enriched at E11.5 in the developing brain derived from aged father. GSEA analysis of differentially up-regulated genes for (a) gene set collection C3 ^45^ and for (b) the NRSF_01 gene set (REST target gene set) in the E11.5 mouse brain derived from aged father.

**Figure S11.**
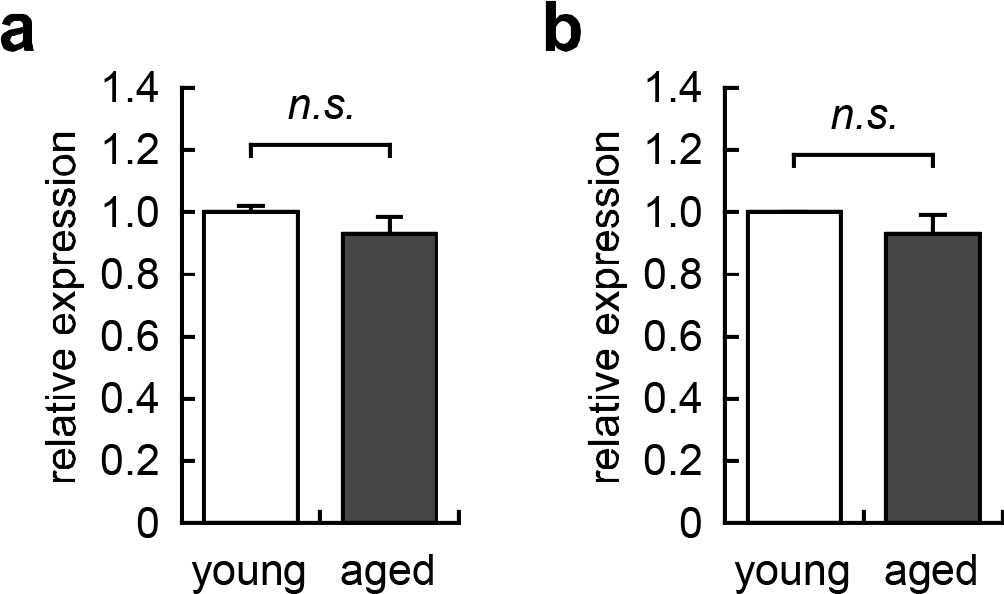
REST/NRSF expression was not changed in offspring brain derived from young and aged fathers. REST/NRSF expression in the offspring brain at (a) E11.5 and (b) E14.5 derived from young and aged fathers in RNA-seq data. The expression level of REST/NRSF is unchanged in the developing brain both at E11.5 and E14.5 of offspring derived from aged father.

## Acknowledgement

This research was supported in part by CREST (to Y.M, T.K, and N.O), the MEXT-Supported Program for the Strategic Research Foundation at Private Universities (to T.K), and JSPS KAKENHI Grants # 25640002, #15K12764 and 16H06530 (to N.O.) and #15K06694 (to K.Y.).

